# FUN-PROSE: A Deep Learning Approach to Predict Condition-Specific Gene Expression in Fungi

**DOI:** 10.1101/2022.06.16.496482

**Authors:** Ananthan Nambiar, Veronika Dubinkina, Simon Liu, Sergei Maslov

## Abstract

mRNA levels of all genes in a genome is a critical piece of information defining the overall state of the cell in a given environmental condition. Being able to reconstruct such condition-specific expression in fungal genomes is particularly important for the task of metabolic engineering of these organisms to produce desired chemicals in industrially scalable conditions. Most of the previous deep learning approaches focused on predicting the average expression levels of a gene based on its promoter sequence, ignoring its variation across different conditions. Here we present FUN-PROSE - a deep learning model trained to predict differential expression of individual genes across various conditions using their promoter sequences and expression levels of all transcription factors. We train and test our model on three fungal species: *Saccharomyces cerevisiae*, *Neurospora crassa* and *Issatchenkia orientalis* and get the correlation between predicted and observed condition-specific gene expression as high as 0.85. We then interpret our model to extract promoter sequence motifs responsible for variable expression of individual genes. We also carried out input feature importance analysis to connect individual transcription factors to their gene targets. A sizeable fraction of both sequence motifs and TF-gene interactions learned by our model agree with previously known biological information, while the rest corresponds to either novel biological facts or indirect correlations.

---

Transcriptional regulation of gene expression is one of the key mechanisms used by biological organisms in general and fungi in particular to modify their phenotype in response to changes in the environment. Protein abundances directly responsible for the phenotypic state of the cell are known to be strongly correlated with mRNA levels of the corresponding genes (for fungi, see e.g.^1–4^). Hence, the ability to predict condition-specific mRNA expression of relevant genes is a crucial step for developing industrial applications of fungal species^5–7^.

Deciphering the complicated process of gene regulation was one of the key research objectives during the past several decades^8–10^. Due to the synchronized work of multiple systems controlling mRNA synthesis and decay^11,12^, the average steady state levels of individual mRNAs may vary from less than one copy per cell to several hundreds per cell across different environmental conditions^13^. The majority of the information shaping this response is encoded in each gene’s two cis-regulatory sequence regions^14^. One of them, referred to as the promoter, is located upstream of the protein coding sequence. It contains sequence motifs recognized by DNA-binding transcription factors (TFs) that enhance or repress mRNA gene expression. Another one is the 3’ UTR sequence located downstream of the protein coding sequence and containing motifs for RNA-binding proteins responsible for mRNA stability and decay^11^. For instance, in the model fungal organism *Saccharomyces cerevisiae*, sequence properties of individual cis-regulatory regions can explain up to half of the variation in mRNA levels across conditions^14^. However, the functional relationship between expression levels of multiple TFs and their gene targets is highly non-linear, and its mechanistic details remain poorly understood, especially in eukaryotic genomes.

Deep neural networks (DNNs) have been extremely successful in learning such complex non-linear relationships in biological data. In particular, convolutional networks (CNNs) are specifically suitable to learn hierarchical patterns in sequence data such as promoters and 3’ UTR regions of individual genes. CNNs were previously applied to extract TF-binding motifs and their higher-order organizational context from ChIP–seq^15,16^, ChIP-exo^17^, and artificial sequence experiments^18^. Other DNN architectures, e.g. fully connected perceptrons, were used to work with other biological data and prediction tasks, e.g., learning the internal state of a cell from gene expression counts^19^. DNNs were also previously used to predict the *average* mRNA level of a gene across many conditions based only on its cis-regulatory sequences^20,21^. Alternatively, there has been work done that incorporates data from transcription factor-DNA binding assays^22^, which is not readily available for many fungal species, to predict expression. However, these studies did not address the question of predicting condition-specific gene expression using sequence information, which is the main subject of our study.

Here we present a broadly applicable DNN model called FUN-PROSE (FUNgal PRomoter to cOndition-Specific Expression) which was trained to predict the relative expression level of a gene in a specific condition based on the gene’s promoter sequence and the expression levels of all TFs of a given fungal species. We tested our model on existing gene expression datasets for three different fungal species and demonstrated its practical applicability not only for model organisms such as *Saccharomyces cerevisiae* but also for less studied fungal species such as *Neurospora crassa* and *Issatchenkia orientalis*, where patterns of gene regulation remain virtually unexplored.

One of the challenges when using DNN models lies in mechanistic interpretation of their results and extraction of new biological knowledge from them^17,23^. To address this challenge, we interrogate our model to identify biologically relevant information of two types. One type is composed of recurrent sequence motifs relevant for regulation of gene expression, e.g., TF-binding motifs. The other type of biological information is the Gene Regulatory Network (GRN) of a species, linking each of the TFs to their gene targets. To learn GRNs in each of our three fungal species, we used input a feature attribution technique to assign tentative TF regulators to individual genes. In *S. cerevisiae*, many sequence motifs and regulatory interactions discovered by our model agree with previously known biological information, while the rest correspond to either novel biological facts or indirect correlations.

In conclusion, our model can be used to both extract new biological knowledge and to tackle a practically important task of manipulating the expression level of a given gene by either changing its promoter sequence or modifying the TF levels.

## RESULTS

To predict the relative expression level of a gene in a particular environmental condition, we reasoned that most of the necessary information should be contained in two sets of data: the promoter sequence of this gene, which contains cis-regulatory sequences recognized by TFs, and individual expression levels of all TFs in this condition. With that in mind, we designed a deep neural network with the following architecture (see Fig. 1A). The first type of inputs (i.e. promoter sequences) is processed through two convolutional layers. The first layer is designed to capture simple sequence motifs in promoters, while the second one should be able to learn combinations of these motifs to account for complex combinatorial effects (e.g. TF-TF interactions, helper proteins, etc.). The second type of inputs (i.e. expression levels of all TFs) is processed through a fully-connected layer. The resulting latent representations are concatenated together and passed through several fully-connected layers that establish a connection between the TFs and corresponding motifs. The final layer then predicts the condition-specific gene expression level. It is important to note that the condition-specific gene expression here is defined as the Z-score of the log-transformed gene expression calculated across all conditions in our data (see Methods). That is, the expression level of each gene is standardized to have the mean of 0 and the standard deviation of 1.

**FIG. 1.**
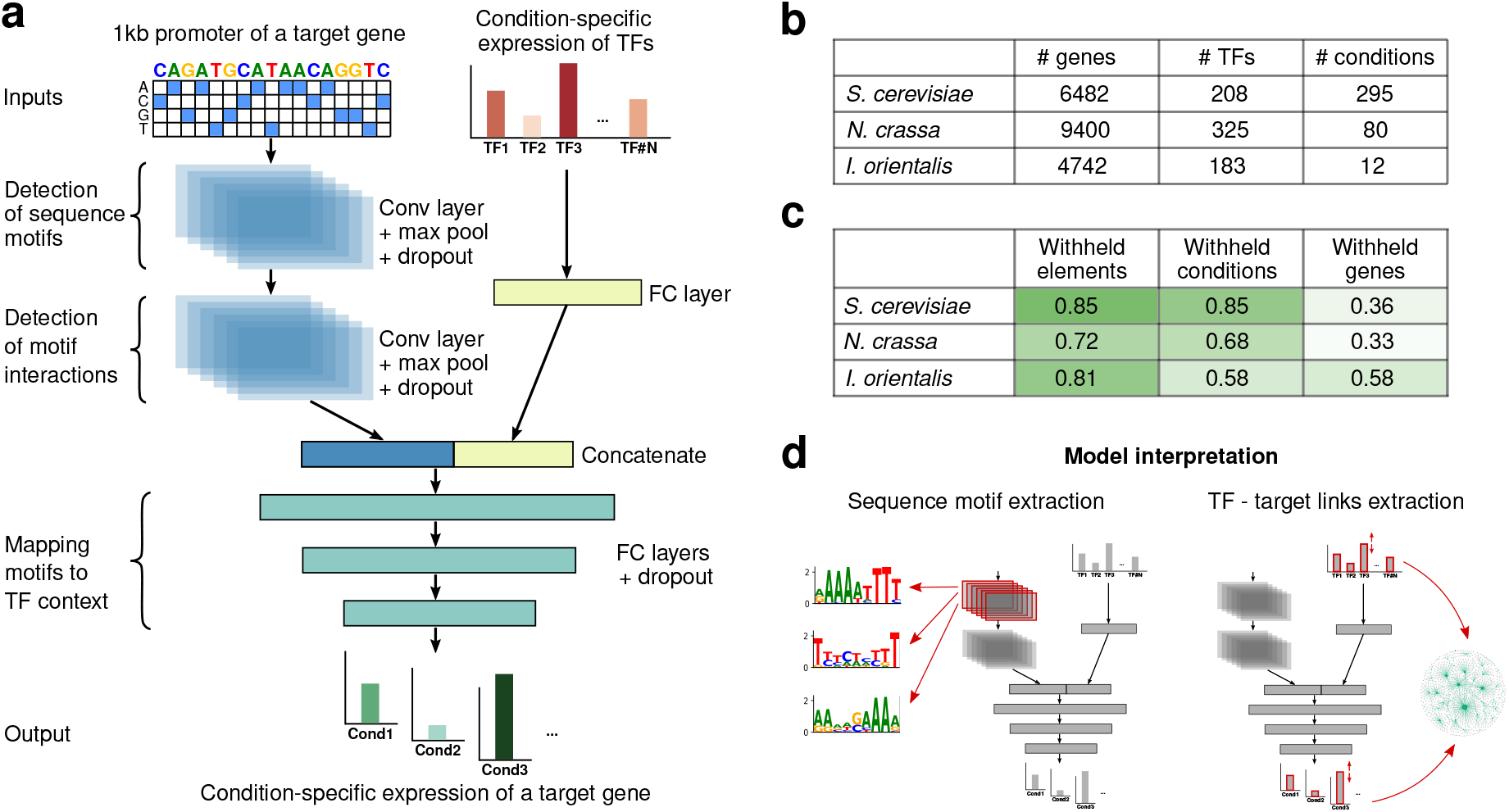
FUN-PROSE model predicts condition-specific gene expression in fungi and allows extracting transcription factor binding motifs and edges of gene regulatory networks. **(a)** Schematic of the FUN-PROSE architecture. The model uses the 1000bp promoter sequence of a gene and condition-specific expression levels of all TFs in the genome as inputs and predicts the expression of this gene in a given condition. FC denotes fully connected layers; Conv denotes convolutional layers. For specific layer parameters (sizes, stride, kernels, etc.), see Table I and Methods. **(b)** The statistics of fungal datasets used in this study. **(c)** FUN-PROSE performance on different datasets for three different train/test set splits (see text for details). **(d)** Schematic of model interpretation procedures to extract sequence motifs (by analysis of the first layer of convolutional filters) and TF-gene regulatory interactions (by Integrated Gradient technique).

To evaluate the performance of FUN-PROSE, we collected several previously published RNA-seq datasets for different fungal species (see Fig. 1B). In particular, we gathered datasets for *Saccharomyces cerevisiae, Neurospora crassa* and *Issatchenkia orientalis* as described in Methods. The compiled *S. cerevisiae* dataset included 6482 genes, 208 TFs and 295 different stress conditions. The *N. crassa* dataset was made up of 9400 genes, 325 TFs and 80 different combinations of growth on different carbon sources and strains with gene knockouts. Finally, the *I. orientalis* dataset was our smallest with 4742 genes, 183 TFs and only 12 conditions of growth on different carbon sources.

The performance of our optimized model architecture for all three species are shown in Fig. 1C. We also took steps to interrogate the network for biologically meaningful information: first is to extract sequence motifs that our DNN model learned during training, we expect some of them to correspond to transcription factor binding motifs; second is to extract TF-gene target links, i.e., edges in GRN using Integrated Gradients^24^ (Fig. 1D).

### Hyperparameter Optimization

In order to make sure that we obtain the best possible performance out of the neural network, we tuned the hyperparameters that define our architecture (see Table I) to maximize the Pearson correlation coefficient of the predicted and true gene expression levels. In our procedure, we used Bayesian optimization in combination with the Async Successive Halving Algorithm (ASHA) scheduler^25^. We performed 250 trials and selected the configuration yielding the highest correlation on the withheld genes data for *N. crassa*. Our hyperparameter search was performed using this dataset because it was our weakest performing result. We split the gene-condition data into train, validation and test sets by randomly withholding 10% of the elements for the validation set and 20% for the test set. The optimal hyperparameters found through this search were then used for all the other species and train/test splits without further hyperparameter tuning due to computational limitations. However, we found that the hyperparameters found by tuning on *N. crassa* worked well for the other two species as well, alluding to the applicability of FUNPROSE across different fungal species.

**TABLE I.** Configuration search space for hyperparameter optimization and best hyperparameters identified in the space. *Uniform* and *LogUniform* indicate that values are uniformly sampled from the domain and log domain of the provided range, respectively. *RandInt* indicates that values are uniformly sampled from integers in the provided range. Bracketed range indicates that values are sampled from the discrete set.

| Hyperparameter Optimizaion Space and Results |  |  |
| --- | --- | --- |
|  | Search Space | Best in Search |
| Promoter Seq. Length | $100 * [2, 3, 4, 5, 6, 7, 8, 9, 10]$ | 1000 |
| Batch Size | [64, 128, 256, 512, 1024] | 256 |
| Peak LR | *LogUniform*[1e-5, 1e-2) | $1.02 * 10^{-4}$ |
| Weight Decay | *LogUniform*[1e-3, 1e-1) | $1.64 * 10^{-3}$ |
| Conv. Layer 1 Kernel Size | [9, 11, 13, 15] | 9 |
| Conv. Layer 1 Kernel # | [16, 32, 64, 128, 256] | 256 |
| Pool. Layer 1 Kernel Size | *RandInt*[5, 20) | 19 |
| Conv. Layer 2 Kernel Size | [9, 11, 13, 15] | 13 |
| Conv. Layer 2 Kernel # | [16, 32, 64, 128, 256] | 64 |
| Pool. Layer 2 Kernel Size | *RandInt*[5, 10) | 8 |
| Conv. Activation Function | [ReLU, GeLU, ELU, SeLU] | ReLU |
| Conv. Dropout | *Uniform*[0, 0.50) | 0.398 |
| TF Hidden Layer Size | [32, 64, 128, 256, 512, 1024] | 1024 |
| TF Hidden Layer Dropout | *Uniform*[0, 0.50) | 0.189 |
| F.C. Activation Function | [ReLU, GeLU, ELU, SeLU] | ELU |
| F.C. Layers Dropout | *Uniform*[0, 0.50) | 0.0166 |

After tuning, our final model’s promoter module was composed of two convolutional layers to process the entire 1kb sequence, where the first layer has 256 filters of 9-bp length to capture relevant sequence motifs, followed by the second layer with 64 filters of length 13 to capture more complex sequence patterns. It is interesting to note that as shown in Table I and Supp. Fig. 4, the optimal kernel size for the first convolutional filter was on the smaller side of the search space, while the pool kernel size for the first layer was on the larger side. This indicates that the first layer of the neural network looks for multiple short motifs and connects them over a longer range.

For the TF-processing fully-connected layer, we found a hidden size of 1024 to be optimal prior to concatenation with the convolution output. We also discovered that applying dropout to the convolutional and fully-connected layers improved our network’s performance.

### Predicting condition-specific gene expression in fungi

For each species, we first split the gene-condition data into train, validation and test sets by randomly withholding 10% of the elements for the validation set and 20% for the test set. That is, when the neural networks are being tested, they will not be receiving the exact combinations of genes and conditions used in training and validation. With this set-up, for *S. cerevisiae*, our neural network achieved a Pearson correlation coefficient of 0.85 between predicted and observed gene expression values. The *N. crassa* and *I. orientalis* models had correlations of 0.72 and 0.81 respectively. This shows that the FUNPROSE framework can be generalized to different fungal species with varying sizes of training datasets.

To further understand the performance of our model, we created scatter plots and confusion matrices for the predictions made on the test set (see Fig. 2). We generated the confusion matrices by trinarizing expression levels on each axis into three sections labelled “Low” (below one standard deviation from the mean), “Mean” (within one standard deviation on either side of the mean) or “High” (above one standard deviation from the mean). The scatter plots and confusion matrices in Fig. 2A-C show that a gene-condition pair predicted to have low expression level is rarely measured to have a high expression level, and vice versa for all three species. Instead, most of the errors seem to arise from genes with either low or high expression levels being predicted to have a mean expression level.

**FIG. 2.**
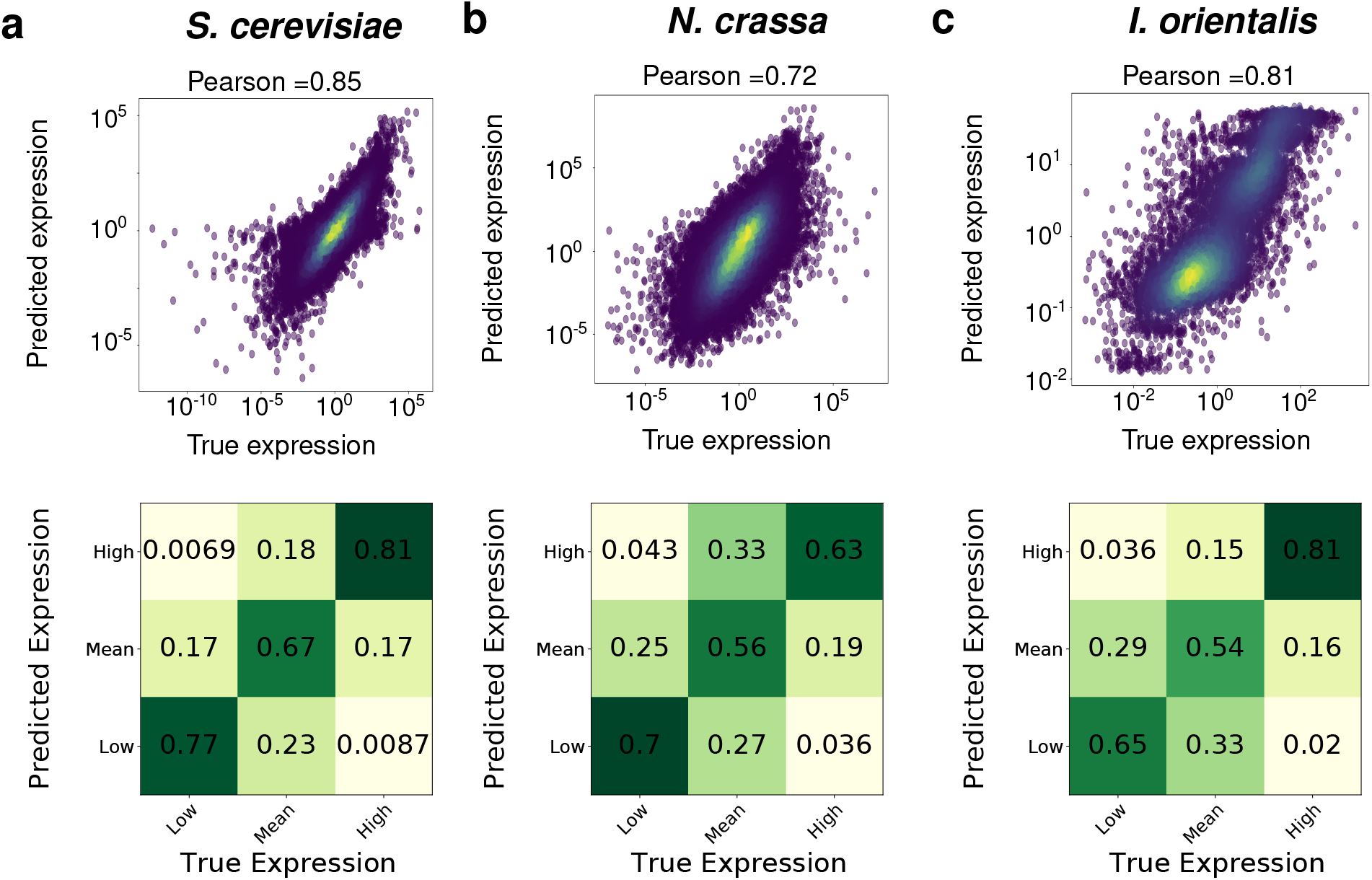
FUN-PROSE model accurately predicts condition-specific gene expression for three different fungal species. The results of FUN-PROSE predicting condition-specific gene expression of *N. crassa* **(a)**, *S. cerevisiae* **(b)**, and *I. orientalis* **(c)**. The top panel shows scatter plots of predicted (y-axis) and experimentally measured (x-axis) expression levels, with the color representing density of points. The bottom panels show confusion matrices of expression levels discretized into three categories (Low, Medium, and High) (see text for details).

We then designed two other more stringent splits to evaluate model performance. One with 10%/20% of conditions withheld for test and validation, i.e, our model is never shown a particular condition at all during training but tries to predict it. This setup allows one to evaluate how well our model will fare in a practical scenario when we use it to predict gene expression in a new condition which has not been experimentally tested as well as the scenario where TF expression levels have been manipulated. In this scenario, the FUN-PROSE model’s performance stayed the same for *S. cerevisiae* and slightly dropped to 0.68 and 0.58 Pearson correlation between predicted and observed gene expression for *N. crassa* and *I. orientalis* respectively. We expect this drop in performance to depend on how different the new, unseen condition is from all other conditions used in the training set.

Another split is to completely withhold some gene promoters during training. This allows to assess how well FUN-PROSE will work for the task of predicting expression of a novel gene in the given condition set. This situation is realized, e.g., if our model is used in evaluating synthetic promoter sequences. This setup is the least accurate with 0.36, 0.33 and 0.58 Pearson correlation between predicted and observed gene expression for *S. cerevisiae*, *N. crassa*, and *I. orientalis* respectively. This drop in performance is to be expected given the fact that our model receives no information about other mechanisms of gene regulation such as epigenetic modification depending on the position of a gene within the genome^26^ and not on its promoter sequence. *I. orientalis* may have better performance than the other two species due to a bimodal distribution of condition-specific gene expression (see Supplementary Figure 3). This bimodal distribution, in turn, may be an artifact of a very small number of conditions in our training set for this species.

### Sequence motifs and their interactions can be extracted from the convolutional filters

We then set to explore the information learned by the FUN-PROSE model trained on each fungal dataset and extract sequence features that were most predictive of gene expression. To do this we used the following procedure: for all genes we extracted all feature maps from the first convolutional layer; then for each of 256 kernels we calculated statistics for base pair frequency in the 9-bp sequence windows around the top-0.5% activations. These sequence motifs quantified by base pair frequency profiles (see Methods) are analogous to Transcription Factor Binding Motifs (TFBMs) traditionally used to quantify sequence patterns recognized by individual TFs. We also calculated positional activation profiles (see Methods) for every extracted sequence motif across all promoters of a given fungal species to look for non-random positional preferences along the promoter sequence.

We hypothesized that sequence motifs extracted from CNN kernels should sometimes match TF-binding motifs. To test this hypothesis, we compared sequence motifs extracted from our model to the known *S. cerevisiae* TFBMs from the YEASTRACT database^27^. We were able to tentatively match 77, 87, and 68 of our 256 sequence motifs to at least one known TFBM in *S. cerevisiae, N. crassa*, and *I. orientalis* genomes respectively (see Supplementary Tables 1-3 and Methods for details). Fig 3A-C shows several examples of motifs extracted from FUN-PROSE model for different species, along with their best match to a known transcription factor binding motif. In the right panel of Fig 3A-C and in Fig 3D we show the positional activation profile of these motifs across all promoter sequences. We found that most sequence motifs extracted from our model exhibit non-random positional preferences indicative of biological function. Indeed, transcriptional regulation typically requires a TF to bind a promoter sequence not too far from the transcription start site^28,29^.

**FIG. 3.**
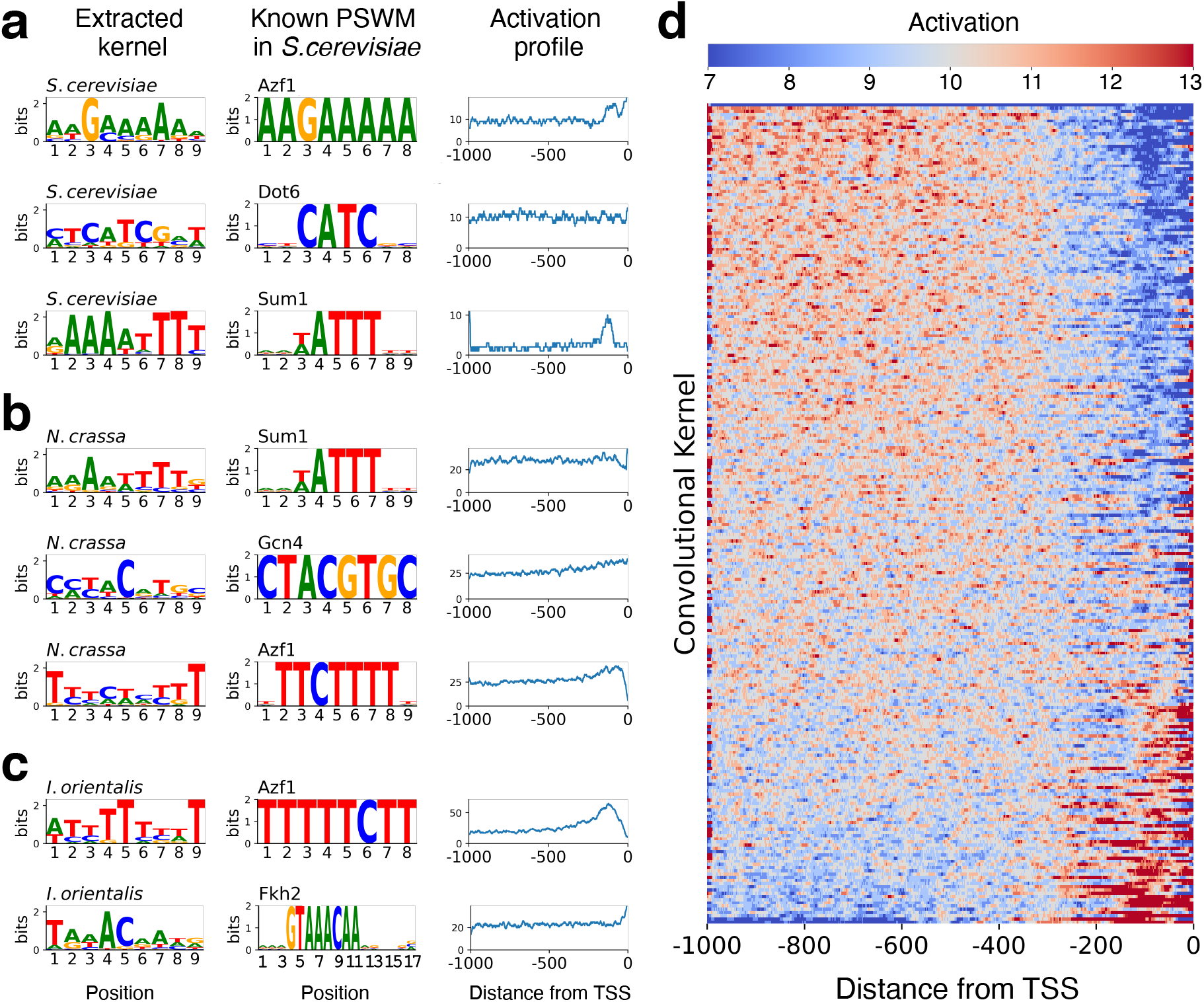
Discovery of sequence motifs by the FUN-PROSE model. **(a)-(c)** Examples of sequence motifs extracted from the convolutional kernels of the FUN-PROSE model trained on the respective species: *S. cerevisiae* **(a)**, *N crassa* **(b)**, *I. orientalis* **(c)**. The best matching *S. cerevisiae* motif in the YEASTRACT database is shown in the middle column and motif’s positional activation profile - in the right column. **(d)** The heatmap showing the positional distribution of the top 0.5% of activations for each motif, i.e. kernel in the first convolutional layer, over all *S. cerevisiae* promoter sequences. The rows of this heatmap are sorted by the average activation level within 300bp from the transcription start site. Note that most motifs exhibit non-random positional preferences indicative of biological function.

Interestingly, for all three species, we independently discovered sequence motifs similar to the one recognized by Azf1 in *S. cerevisiae.* Azf1 is known to be a transcription activator of genes involved in carbon metabolism and energy production^30^ and is expected to be actively working in the set of conditions we used for our model training for all three species.

### Regulatory interactions between transcription factors and target genes can be inferred from the neural network

Before trying to understand which TFs are the most predictive ones for condition-specific expression of a particular gene, we verified that expression levels of individual TFs are indeed used by the FUN-PROSE model to make predictions. We trained three new models for each of our fungal species where, where instead of sending the expression levels of individual TFs as inputs, we send only one given by the mean expression level across all TFs. This singular value is then concatenated to the outputs of the convolutional layers before being fed to the fully connected layers to predict condition-specific gene expression. When this network was evaluated on the withheld elements datasets, we obtained correlation coefficients of 0.36, 0.34 and 0.55 for *S. cerevisiae, N. crassa* and *I. orientalis*, respectively. These are much lower than the correlations of 0.85, 0.72 and 0.81 obtained by our original FUN-PROSE model. This alleviates a potential concern that only the overall levels of TF expression for a cell are necessary to make predictions. Instead, this shows that the expression levels of individual TFs play an important role in accurately predicting condition-specific gene expression.

To understand the role of different TFs in making predictions, we generated a list of TF-target gene interactions for the model trained on the *S. cerevisiae* dataset using Integrated Gradients technique^24^, which quantifies how much each of the input variables contributed to the final prediction of a given output (see Methods). This data was then binarized using the threshold of 3 standard deviations from the mean absolute TF-gene score to create a network of TF-target gene interactions made up of 21 TFs, 1392 target genes and 4756 edges as shown in Fig 4A. The size of the nodes and their labels represent the out-degree of a given TF. As seen in this network diagram, a handful hub TFs that make up the most of TF-gene interactions. The shape of the cumulative histogram of out-degrees Fig 4B shows a sharp transition between around 10 TF hubs and the rest of TFs. In fact, these 10 hubs cover 95.7% of edges in the network. These results indicate that, when considering stress response in *S. cerevisiae*, where our training data came from, a few TFs are sufficiently predictive of condition-specific gene expression. The reasons behind the predictive power of these specific TFs is better understood by taking a closer at their biological function. The top five TFs by out-degree are DOT6, MBF1, MATALPHA1, HMRA1 and GCN4 included in Fig 4D. DOT6 has been previously shown to be a master regulator of the stress response^31^. Going down the list, MBF1 and GCN4 TFs were also shown to be potentially interacting master regulators in response to stress in yeast, especially nutritional stress^32,33^.

**FIG. 4.**
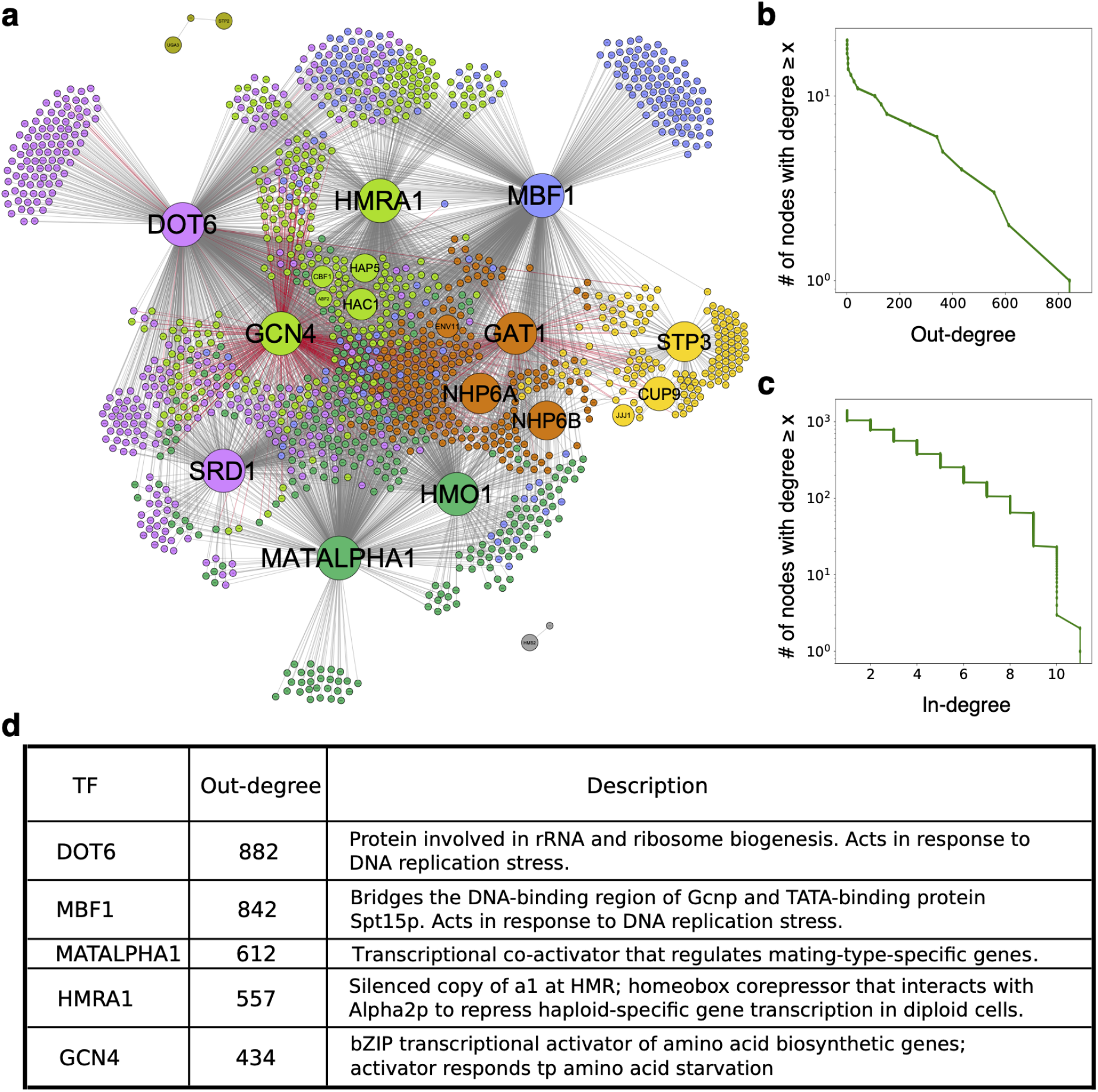
*S. cerevisiae* Gene Regulatory Network learned by the FUN-PROSE model. **(a)** The network of TF-target gene interactions obtained by applying a 3 standard deviation threshold to the TF-target gene Integrated Gradients scores for the *S. cerevisiae* dataset. Red edges mark experimentally confirmed interactions from the YEASTRACT database. Nodes are colored by clusters obtained by modularity optimization. **(b)** Cumulative histogram (number of nodes with degree >= *x*) of out-degrees of TFs and **(c)** in-degrees of target genes. **(d)** A table of properties of the top TFs with the highest degree-centrality, including their out-degree and the biological function according to the Saccharomyces Genome Database (SGD).

Fig 4C shows that the in-degree of gene targets (the number of TFs regulating a given gene) approximately follows an exponential distribution, This is consistent with previous results obtained for experimentally observed regulatory interactions in *S. cerevisiae*^34^.

While only a small proportion of links in Fig 4A have experimental validation (red links), it is important to note that the links we discover are not necessarily direct causal interactions. This is because machine learning methods routinely pick up indirect interactions, such as when a TF may regulate the expression of another (hidden) TF, which in-turn regulates the expression of a target gene. However, an interesting aspect about our inferred network is that the two TFs with the highest proportion of experimentally confirmed links are GCN4 and GAT1. Both of them are well known stress response regulators, with GCN4 mentioned above and GAT1 being linked to salt stress response.

Next, the network was analyzed by clustering the nodes to maximize the modularity of the network (shown as node color on Fig 4 A)^35^. Then, we ran Gene Ontology (GO) term enrichment analysis on genes in each cluster^36^. Through this, we saw that often, the GO terms associated with a module could be explained by the TFs in the cluster. For example, most of the GO terms associated with the Yellow cluster were related to RNA processing and ribosome biogenesis. The TFs in the cluster include JJJ1 which has been shown to be involved in 60S ribosomal subunit biogenesis^37^. Another TF in the Yellow cluster is STP3, a protein similar to STP1. While STP3 has not been widely studied, STP1 is known to be involved in tRNA splicing^38^. Next, the Blue cluster had GO terms mostly associated with ion transport. This aligns with evidence showing how MBF1 is associated with iron transport in another species of yeast, *C. albicans*, indicating that a similar association might also be present in *S. cerevisiae*^32^. Another example of how the GO term of a cluster matches what is known about the TFs that regulate the cluster is the Brown cluster, which is mostly assigned GO terms relating to DNA repair. As it turns out, NHP6A/B loss leads to increased genomic instability, hypersensitivity to DNA-damaging agents^39^.

## DISCUSSION

The focus of this study is on prediction of condition-specific gene expression in fungi. This prediction task is unlike most previous attempts at predicting gene expression^20,21^ because we aim to predict the variation of expression of each gene across different conditions (Z-score), instead of predicting its absolute expression level (mRNAs/cell) averaged over all conditions. To do so, we use as inputs, promoter sequences (to encode the information about genes) and the expression levels of all TFs (to encode the information about conditions). These inputs are processed by the FUN-PROSE network, which is made up of a convolutional neural network to extract features from the promoter sequences, a fully-connected feed-forward network to learn features from the TF expression levels.

We found that it is possible to predict condition-specific gene expression with high accuracy for three different fungal species: *S. cerevisiae*, *N. crassa* and *I. orientalis*, indicating that FUN-PROSE accuracy generalizes well. We also found that FUN-PROSE model can also be applied to predict gene expression in previously unseen conditions and, separately, on previously unseen promoter sequences, although the prediction accuracy on new promoter sequences was considerably lower than that for previously unseen conditions. These results indicate that our model could have practical applications in predicting how the transcriptome of a fungal species will react to a new condition that has not been tested yet, predicting the consequences of TF knockout/overexpression as well as predicting the effect of promoter modification.

Next, we showed that FUN-PROSE can be successfully interpreted to extract the biological information it used to make predictions. We did this in two parts: first, we interpreted the convolutional neural network module of the FUN-PROSE model to extract the sequence motifs. Second, we interpreted the fully-connected module to extract interactions between TFs and their gene targets. This was done using Integrated Gradients, allowing one to predict which inputs significantly contributed to prediction of a given output variable (gene expression). The results of both exercises were compared to the existing biological knowledge, which is especially significant for *S. cerevisiae*. The fact that a sizeable fraction of our predictions overlapped with known TFBS or regulatory interactions convinced us that the FUN-PROSE model uses biologically relevant information to make its predictions. This also alludes to how we might be able to use FUN-PROSE to generate novel biological hypotheses for less studied species.

In order to understand if additional information regarding cis-regulatory sequences of genes would help our model, we trained a variant of our model with an additional input of 1000bp from 3’ UTR located downstream from the protein coding region of the gene. Similar to the promoter sequence, this information was processed by a two-layer CNN. The latent representations of the 3’ UTR is then concatenated with that of the promoter and the TF expression levels before being fed to the fully connected layers. With this additional piece of input, for the withheld elements split of the data, the model was able to achieve correlation coefficients of 0.87, 0.77 and 0.83 in *S. cerevisiae*, *N. crassa* and *I. orientalis*, respectively. The performance of this model is somewhat higher than that of the original FUN-PROSE model: 0.85, 0.72 and 0.81. We expected that adding 3’ UTR region controlling mRNA degradation would lead to an improvement in accuracy of our predictions. The fact that the magnitude of this improvement was relatively small could be tentatively attributed to the fact that we did not explicitly include condition-specific expression levels of proteins controlling mRNA degradation. So, the model had to use indirect relationships due to co-expression of these proteins with transcription factors. In other words, TF expression levels impact the expression levels of RNA binding proteins, which in turn affect the mRNA decay. Future work could extend our model by identifying families of proteins responsible for mRNA degradation and including their expression levels as inputs alongside TFs.

Many sequence motifs we discovered for different species are similar to each other. In order to quantify the overall level of conservation of these motifs, we performed the following experiment: we took the weights of the convolutional layers from the *S.cerevisiae* model, froze them and retrained the rest of the model for *I. orientalis*. The final model performance for withheld genes test/train split was Pearson correlation equal to 0.45, which is still significant but somewhat smaller than Spearman correlation equal to 0.58 in the FUN-PROSE model in which CNN weights were independently trained.

The success of FUN-PROSE across different species in the fungal kingdom suggests that it might be generalizable to other kingdoms, e.g. plants and animals. In doing so, one of the potential modifications of the model is changing the length of the promoter sequence that the model takes. The expression of some genes in metazoan species is known to be regulated by distal transcriptional enhancers^40^. Directly incorporating these distal enhancers in our modeling framework may not be computationally tractable. One possible solution to this problem is to extend our model by taking into account the 3D chromosome structure connecting enhancers to their gene targets. One can also incorporate into our model additional input features that might be especially important for more complex organisms. One example of this is the information about epigenetic modifications in the neighborhood of a given gene.

Another possible direction for a future study is to predict condition-specific gene expression using single cell transcriptomics data. Machine learning models may potentially perform better in this setting than for spatially averaged expression data due to the lack of averaging over distinct subpopulations.

## METHODS

### Data sources and preprocessing

We used previously published RNA-seq data on *N. crassa* (wild type and gene-deletion mutants) growing on different carbon sources^41^, recent *S. cerevisiae* RNA-seq data for 28 analog sensitive kinase alleles across 12 different conditions (stresses and different media) (GEO: GSE115556)^42^ and *I. orientalis* RNA-sec data for growth in different media conditions (YPD+glucose and lignocellulosic extracts).

#### Processing the expression data

The data was processed and raw/FPKM counts were obtained by the authors of the respective studies. To standardize the data, for each media condition we first renormalized raw counts to FPKM (if not already) and averaged data for multiple replicates of the same condition. We then filter out genes that have mean expression below 0.05 and genes that have coefficient of variation below 0.3. Supplementary Figures 1-3 (top-left) shows the distribution of counts after applying the two filters. Finally, we log-transformed counts and performed z-score normalization for each gene. The result of these transformations on the distribution of counts is shown in Supplementary Figures 1-3 (top-right).

#### Processing the sequence data

For *N. crassa* we used reference genome Neurospora crassa OR74A v2.0^43^ obtained from MycoCosm; for *S. cerevisiae* we used S. cerevisiae S288C R64-3-1^44^ obtained from SGD; and for *I. orientalis* we used Pichia kudriavzevii CBS573^45^ obtained from MycoCosm. For each gene we defined promoter sequence as 1kb upstream of the start codon and extracted them from the corresponding reference genomes. We filtered out genes that had promotersshorter than 1kb as it sometimes happens on the ends of chromosome. In the end, we worked with 9725, 6645, and 4925 genes for which we had all 3 types of data for *N. crassa*, *S. cerevisiae*, and *I. orientalis* respectively.

#### Predicting transcription factors

We used InterProScan v5.52-86.0 to annotate reference genomes. To obtain all putative TFs for each species, we used this annotation to extract genes that correspond to the list of TF-specific pfams from DNA-binding do main database (DBD)^46^ v2.03 and TF-specific Interpro terms from Fungal Transcription Factor Database (FTFD)^47^ v1.2. Overall we identified 325, 208, and 183 putative TFs in *N. crassa*, *S. cerevisiae*, and *I. orientalis* genomes respectively.

### The model

As shown in Figure 1, our final neural network is composed of three modules: a convolutional neural network that takes the promoter sequence as input, a feed-forward layer that takes the transcription factor expression levels as input, and a multi-layer feed-forward neural network that takes the concatenation of the outputs of the previous two modules as input.

The convolutional neural network module is composed of two convolutional layers. The first convolutional layer has 256 kernels of length 9 and stride of one. This is followed by max pooling with kernel size and stride of 19, a ReLU activation function, dropout and batch normalization. The second convolutional layer, on the other hand, has 64 kernels. Each of these kernels has a length of 13 and a stride of one. This layer is followed by max pooling with kernel size and stride of 8, a ReLU activation function, dropout and batch normalization.

The feed-forward layer that processes the TF expression levels is a fully-connected layer with a ELU activation function.

Finally, the outputs of the convolutional module and the feed-forward module are concatenated and sent through a final module with three fully-connected layers with ELU activation, dropout, batch normalization, and then a final fully-connected layer to predict the expression of a particular gene.

### Hyperparameter Optimization and Model Training

During training, we defined the loss function as the mean squared error of the gene expression level predictions. The AdamW optimizer was used to minimize this loss. To select the optimal set of hyperparameters, we looked at the configuration yielding the highest validation Pearson correlation.

During hyperparameter optimization, we allowed a maximum of 20 epochs for each trial, with a minimum of 5 epochs before stopping. Trials were also stopped early if they reached a plateau, as defined by the standard deviation of the validation correlation coefficient not exceeding 0.01 in the final 5 epochs. The ASHA scheduler was configured with a reduction factor of 3 and 1 bracket. We used the following software for hyperparameter optimization: Ray v1.8.0, PyTorch v1.9.1, and CUDA v11.5. Models were trained on an NVIDIA V100 GPU with 16GB of RAM using automatic mixed-precision training.

In model training, we allowed for a maximum of 60 epochs and training was stopped early if the validation correlation coefficient did not improve for 5 epochs in a row. We ran our model training on a NVIDIA GeForce GTX 1080 Ti GPU.

### Interpreting the convolutional kernels

#### Inferring promoter motifs from convolutional kernels

We inferred promoter motifs learned by each trained model by examining the 256 kernels in the first convolutional layer, which capture such information^23^. For each kernel *x*, denoted by *Conv1d_x_*, we generated a feature map Fx of dimension *N* × 5 × 1000 as the output of processing all N unique one-hot-encoded 1000-bp promoter sequences (*P*_1_,…, *P_N_*). For notation, the xth feature map *F_x_* is indexed as 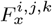 where *i* identifies the promoter, *j* identifies the nucleotide base (A, C, G, T, and N to represent an unknown base), and *k* identifies the sequence position. The *i*th promoter *P_i_* is indexed as 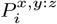 where *x* identifies the nucleotide base and *y*: *z* identifies an inclusive position range. So then:

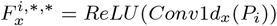

We then constructed a motif representation *T_x_* with dimension 5×9 for each kernel from its feature map *F_x_* as a weighted aggregate of the 9-bp sequence windows corresponding to the top 0.5% activations in *F_x_* (denoted by the pair of promoter and position index lists 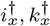). *T_x_* is indexed as 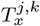 where *j* identifies the nucleotide base and *k* identifies a position. *Mx* is indexed similarly.

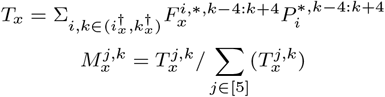

The final representation *M_x_* is a position-specific weight matrix (PSWM) for the *x*th kernel.

#### Positional activation profiles for convolutional kernels

We also constructed positional activation profiles for each kernel as the counts of the top 0.5% of non-zero activations over the promoter sequences. Values were smoothed using a moving average over the sequence with a window size of 15 bps.

#### Comparing against previously reported motifs

We compared these motifs against reported TF-binding *S. cerevisiae* motifs in the YEASTRACT database (version 20130918)^27^ using *Tomtom.*^48^. For motif comparison, we used the Pearson Correlation Coefficient function and complete scoring with a statistical significance threshold of E-value < 0.5.

### Interpreting TF-gene relationships

#### Generating a table of TF-gene interactions

To investigate the importance of TFs for prediction, we used Integrated Gradient (IG) scores^24^. For each prediction made in the withheld elements test set, we calculated IG scores using Captum v0.5.0 with zero-tensors as the baselines and 20 approximation steps. We only used the IG scores for TF expression levels and averaged them for each target gene. This gives us a table of TF-gene interactions in the model. To find the overall importance of each TF, we then averaged the values of this table for each TF across all target genes.

#### Creating and analyzing a network of TF-target gene interactions

The table of TF-target gene relationships was then thresholded (3 standard deviations from the mean for *S. cerevisiae*) to obtain a network of TF-gene relationships. This network was vizualized using Gephi^49^.

To perform our GO enrichment analysis, we first clustered our network by maximizing modularity, a measure of the density of links inside communities as compared to links between communities^35^. Once again, we did this using a built in function in Gephi^49^. Using the clusters obtained, we then performed Gene Ontology (GO) term enrichment analysis to identify terms that are significantly overrepresented in each cluster. This was conducted with the BiNGO tool using a hypergeometric statistical test and a Benjamini-Hochberg FDR corrected significance level of α = 0.05^36^.

## Code availability

All code for simulations used in this manuscript can be found at https://github.com/maslov-group/FUN-PROSE.

## Acknowledgements

This work was funded by the DOE Center for Advanced Bioenergy and Bioproducts Innovation (U.S. Department of Energy, Office of Science, Office of Biological and Environmental Research under Award Number DE-SC0018420). Any opinions, findings, and conclusions or recommendations expressed in this publication are those of the authors and do not necessarily reflect the views of the U.S. Department of Energy.

Part of this work was performed under the auspices of the U.S. Department of Energy by Argonne National Laboratory under Contract DE-AC02-06-CH11357.

This work utilizes resources supported by the National Science Foundation’s Major Research Instrumentation program, grant #1725729, as well as the University of Illinois at Urbana- Champaign^50^.

S.L. has been supported by the James Scholar Honors Program and the Illinois Scholars Undergraduate Research Program.

We thank Peter Koo and Saurabh Sinha for insightful discussions.

## Author contributions

All authors designed the study. S.M. supervised the study; A.N. and S.L. performed simulations and calculations. V.D. curated the data; All authors discussed and wrote the paper.

## References

1 S. P. Gygi, Y. Rochon, B. R. Franza, and R. Aebersold, Molecular and cellular biology 19, 1720 (1999).

2 D. Greenbaum, C. Colangelo, K. Williams, and M. Gerstein, Genome biology 4, 1 (2003).

3 T. J. Griffin, S. P. Gygi, T. Ideker, B. Rist, J. Eng, L. Hood, and R. Aebersold, Molecular & Cellular Proteomics 1, 323 (2002).

4 Y. Liu, A. Beyer, and R. Aebersold, Cell 165, 535 (2016).

5 D. E. Cameron, C. J. Bashor, and J. J. Collins, Nature Reviews Microbiology 12, 381 (2014).

6 D. G. Michael, E. J. Maier, H. Brown, S. R. Gish, C. Fiore, R. H. Brown, and M. R. Brent, Proceedings of the National Academy of Sciences 113, E7428 (2016).

7 J. Nielsen and J. D. Keasling, Cell 164, 1185 (2016).

8 P. Kemmeren, K. Sameith, L. A. Van De Pasch, J. J. Benschop, T. L. Lenstra, T. Margaritis, E. O’Duibhir, E. Apweiler, S. van Wageningen, C. W. Ko, et al., Cell 157, 740 (2014).

9 S. R. Hackett, E. A. Baltz, M. Coram, B. J. Wranik, G. Kim, A. Baker, M. Fan, D. G. Hendrickson, M. Berndl, and R. S. McIsaac, Molecular systems biology 16, e9174 (2020).

10 Y. Kang, N. R. Patel, C. Shively, P. S. Recio, X. Chen, B. J. Wranik, G. Kim, R. S. McIsaac, R. Mitra, and M. R. Brent, Genome research 30, 459 (2020).

11 B. C. Foat, S. S. Houshmandi, W. M. Olivas, and H. J. Bussemaker, Proceedings of the National Academy of Sciences 102, 17675 (2005).

12 S. Inukai, K. H. Kock, and M. L. Bulyk, Current opinion in genetics & development 43, 110 (2017).

13 L. Wodicka, H. Dong, M. Mittmann, M.-H. Ho, and D. J. Lockhart, Nature biotechnology 15, 1359 (1997).

14 J. Cheng, K. C. Maier, Ž. Avsec, P. Rus, and J. Gagneur, Rna 23, 1648 (2017).

15 J. Yang, A. Ma, A. D. Hoppe, C. Wang, Y. Li, C. Zhang, Y. Wang, B. Liu, and Q. Ma, Nucleic acids research 47, 7809 (2019).

16 D. Quang and X. Xie, Methods 166, 40 (2019).

17 Ž. Avsec, M. Weilert, A. Shrikumar, S. Krueger, A. Alexandari, K. Dalal, R. Fropf, C. McAnany, J. Gagneur, A. Kundaje, et al., Nature Genetics 53, 354 (2021).

18 E. D. Vaishnav, C. G. de Boer, J. Molinet, M. Yassour, L. Fan, X. Adiconis, D. A. Thompson, J. Z. Levin, F. A. Cubillos, and A. Regev, Nature 603, 455 (2022).

19 C. Culley, S. Vijayakumar, G. Zampieri, and C. Angione, Proceedings of the National Academy of Sciences 117, 18869 (2020).

20 V. Agarwal and J. Shendure, Cell reports 31, 107663 (2020).

21 J. Zrimec, C. S. Börlin, F. Buric, A. S. Muhammad, R. Chen, V. Siewers, V. Verendel, J. Nielsen, M. Töpel, and A. Zelezniak, Nature communications 11, 1 (2020).

22 Q. Song, J. Lee, S. Akter, M. Rogers, R. Grene, and S. Li, Nucleic Acids Research 48, e62 (2020).

23 P. K. Koo and M. Ploenzke, Nature Machine Intelligence 3, 258 (2021).

24 M. Sundararajan, A. Taly, and Q. Yan, in Proceedings of the 34th International Conference on Machine Learning - Volume 70, ICML’17 (JMLR.org, 2017) p. 3319–3328.

25 L. Li, K. Jamieson, A. Rostamizadeh, E. Gonina, J. Bentzur, M. Hardt, B. Recht, and A. Talwalkar, in Proceedings of Machine Learning and Systems, Vol. 2, edited by I. Dhillon, D. Papailiopoulos, and V. Sze (2020) pp. 230–246.

26 S. I. Grewal and D. Moazed, science 301, 798 (2003).

27 M. C. Teixeira, P. T. Monteiro, J. F. Guerreiro, J. P. Gonçalves, N. P. Mira, S. C. dos Santos, T. R. Cabrito, M. Palma, C. Costa, A. P. Francisco, S. C. Madeira, A. L. Oliveira, A. T. Freitas, and I. Sá-Correia, Nucleic Acids Research 42, D161 (2013), https://academic.oup.com/nar/article-pdf/42/D1/D161/3525787/gkt1015.pdf.

28 I. Erb, and E. Van Nimwegen, PLoS One 6, e24279 (2011).

29 M. J. Rossi, P. K. Kuntala, W. K. Lai, N. Yamada, N. Badjatia, C. Mittal, G. Kuzu, K. Bocklund, N. P. Farrell, T. R. Blanda, et al., Nature 592, 309 (2021).

30 M. G. Slattery, D. Liko, and W. Heideman, Eukaryotic cell 5, 313 (2006).

31 A. A. Granados, J. M. J. Pietsch, S. A. Cepeda-Humerez, I. L. Farquhar, G. Tkačik, and P. S. Swain, Proceedings of the National Academy of Sciences 115, 6088 (2018), https://www.pnas.org/doi/pdf/10.1073/pnas.1716659115.

32 S. Amorim-Vaz, A. T. Coste, V. D. T. Tran, M. Pagni, and D. Sanglard, Frontiers in Fungal Biology 2 (2021), 10.3389/ffunb.2021.658899.

33 A. G. Hinnebusch and K. Natarajan, Eukaryotic Cell 1, 22 (2002), https://journals.asm.org/doi/pdf/10.1128/EC.01.1.22-32.2002.

34 S. Maslov and K. Sneppen, Physical biology 2, S94 (2005).

35 V. D. Blondel, J.-L. Guillaume, R. Lambiotte, and E. Lefebvre, Journal of Statistical Mechanics: Theory and Experiment 2008, P10008 (2008).

36 S. Maere, K. Heymans, and M. Kuiper, Bioinformatics 21, 3448 (2005).

37 A. E. Meyer, N.-J. Hung, P. Yang, A. W. Johnson, and E. A. Craig, Proceedings of the National Academy of Sciences 104, 1558 (2007), https://www.pnas.org/doi/pdf/10.1073/pnas.0610704104.

38 M. U. Jørgensen, C. Gjermansen, H. A. Andersen, and M. C. Kielland-Brandt, Current Genetics 31, 241 (1997).

39 S. Giavara, E. Kosmidou, M. Hande, M. E. Bianchi, A. Morgan, F. d’Adda di Fagagna, and S. P. Jackson, Current Biology 15, 68 (2005).

40 M. Bulger and M. Groudine, Cell 144, 327 (2011).

41 V. W. Wu, N. Thieme, L. B. Huberman, A. Dietschmann, D. J. Kowbel, J. Lee, S. Calhoun, V. R. Singan, A. Lipzen, Y. Xiong, et al., Proceedings of the National Academy of Sciences 117, 6003 (2020).

42 K. Mace, J. Krakowiak, H. El-Samad, and D. Pincus, PloS one 15, e0230246 (2020).

43 J. E. Galagan, S. E. Calvo, K. A. Borkovich, E. U. Selker, N. D. Read, D. Jaffe, W. FitzHugh, L.-J. Ma, S. Smirnov, S. Purcell, et al., Nature 422, 859 (2003).

44 S. R. Engel, F. S. Dietrich, D. G. Fisk, G. Binkley, R. Balakrishnan, M. C. Costanzo, S. S. Dwight, B. C. Hitz, K. Karra, R. S. Nash, S. Weng, E. D. Wong, P. Lloyd, M. S. Skrzypek, S. R. Miyasato, M. Simison, and J. M. Cherry, G3 Genes—Genomes—Genetics 4, 389 (2014), https://academic.oup.com/g3journal/article-pdf/4/3/389/37120100/g3journal0389.pdf.

45 A. P. Douglass, B. Offei, S. Braun-Galleani, A. Y. Coughlan, A. A. Martos, R. A. Ortiz-Merino, K. P. Byrne, and K. H. Wolfe, PLoS pathogens 14, e1007138 (2018).

46 D. Wilson, V. Charoensawan, S. K. Kummerfeld, and S. A. Teichmann, Nucleic acids research 36, D88 (2008).

47 J. Park, J. Park, S. Jang, S. Kim, S. Kong, J. Choi, K. Ahn, J. Kim, S. Lee, S. Kim, et al., Bioinformatics 24, 1024 (2008).

48 S. Gupta, J. A. Stamatoyannopoulos, T. L. Bailey, and W. S. Noble, Genome Biology 8, R24 (2007).

49 M. Bastian, S. Heymann, and M. Jacomy, “Gephi: An open source software for exploring and manipulating networks,” (2009).

50 V. Kindratenko, D. Mu, Y. Zhan, J. Maloney, S. H. Hashemi, B. Rabe, K. Xu, R. Campbell, J. Peng, and W. Gropp, “Hal: Computer system for scalable deep learning,” in Practice and Experience in Advanced Research Computing (Association for Computing Machinery, New York, NY, USA, 2020) p. 41–48.

