## Supplementary Figures for "FUN-PROSE: A Deep Learning Approach to Predict Condition-Specific Gene Expression in Fungi"

### Supplementary Information

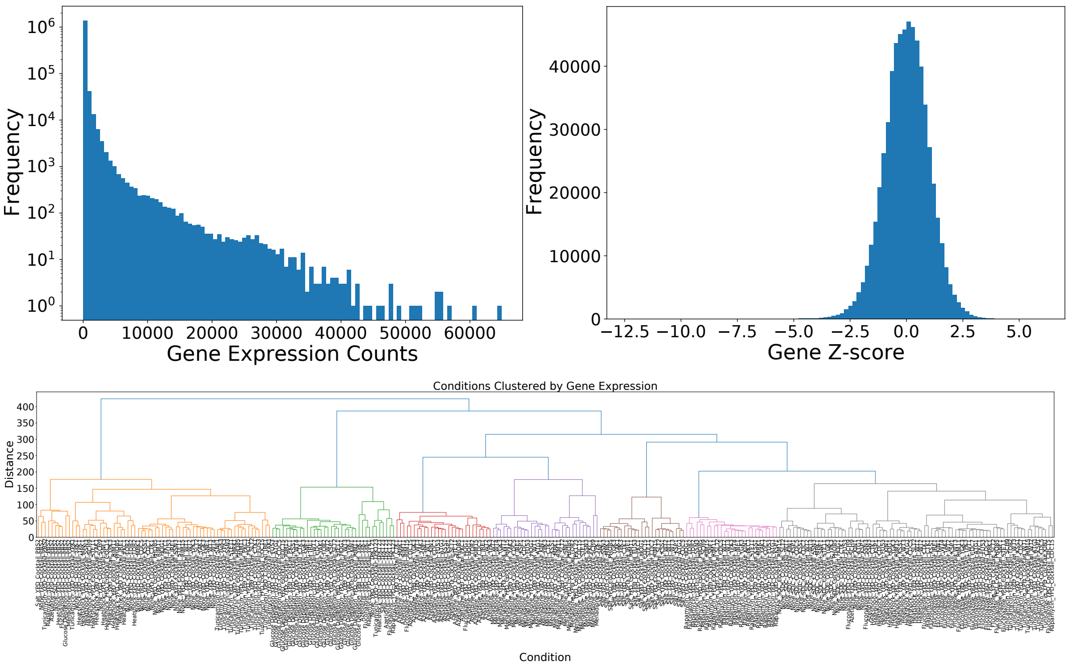


SUPP.FIG. 1. **Vizualizing the *Saccharomyces cerevisiae* dataset. (Top-left)** Histogram of gene expression with CV filter. **(Top-right)** Histogram of Z-scored expressions. **(Bottom)** The different conditions in the dataset clustered by gene expression using agglomerative clustering.

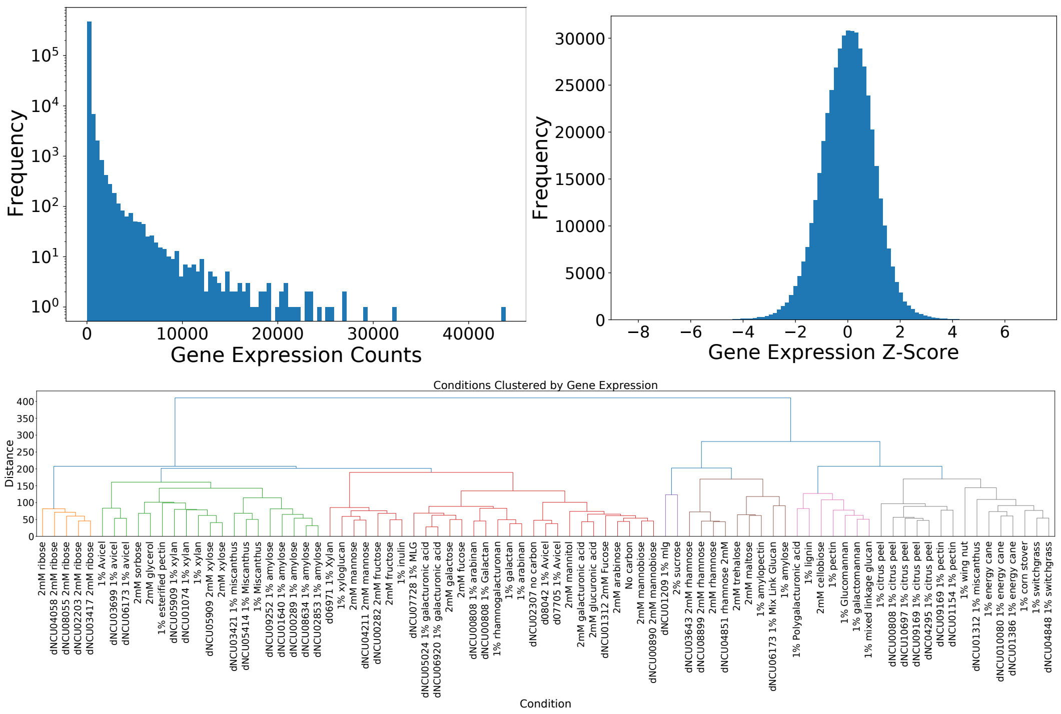


SUPP.FIG. 2. **Vizualizing the *Neurospora crassa* dataset. (Top-left)** Histogram of gene expression with CV filter. **(Top-right)** Histogram of Z-scored expressions. **(Bottom)** The different conditions in the dataset clustered by gene expression using agglomerative clustering.

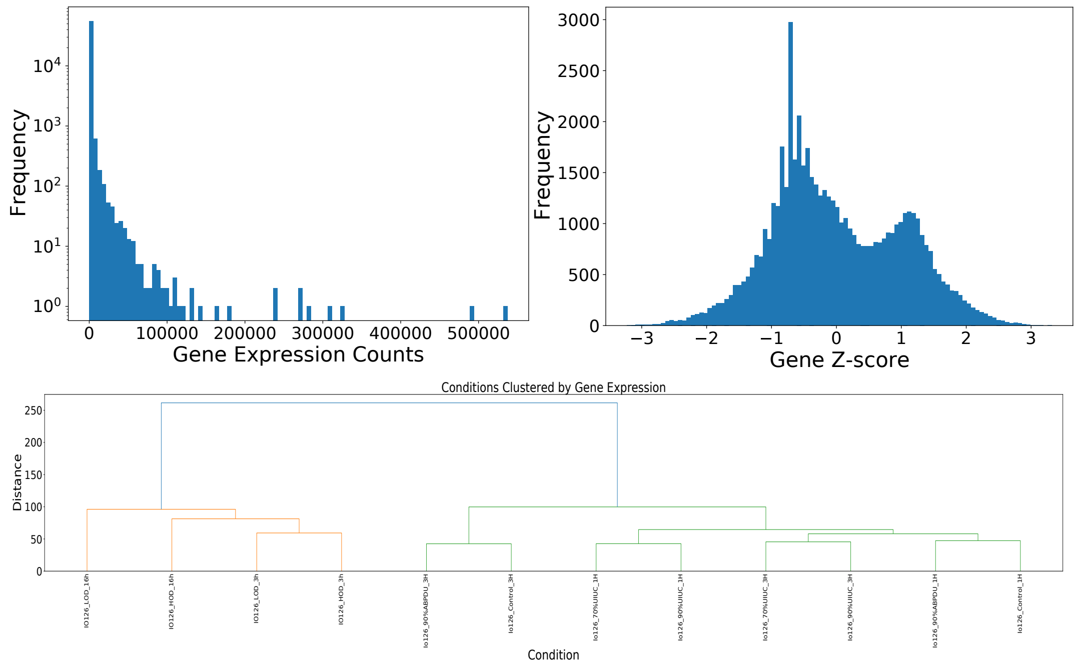


SUPP.FIG. 3. **Vizualizing the *Issatchenkia orientalis* dataset.** **(Top-left)** Histogram of gene expression with CV filter. **(Top-right)** Histogram of Z-scored expressions. **(Bottom)** The different conditions in the dataset clustered by gene expression using agglomerative clustering.

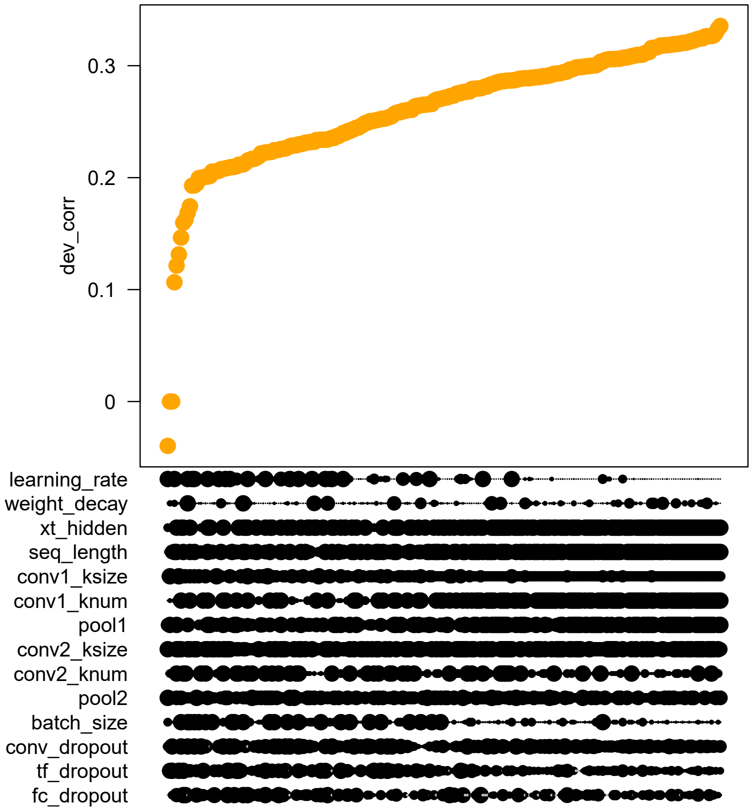


SUPP.FIG. 4. **The factor-response plot for our hyperparameter optimization runs.** The y-axis of the plot shows the correlation between predicted and measured expression for the validation set. The trials on the plot are sorted in ascending correlations. On the x-axis, we show the various hyperparameters that we are optimizing. The size of the black markers represent the value of the hyperparameter for that trial. For example, this plot can be interpreted to show that a smaller learning rate leads to better performance.

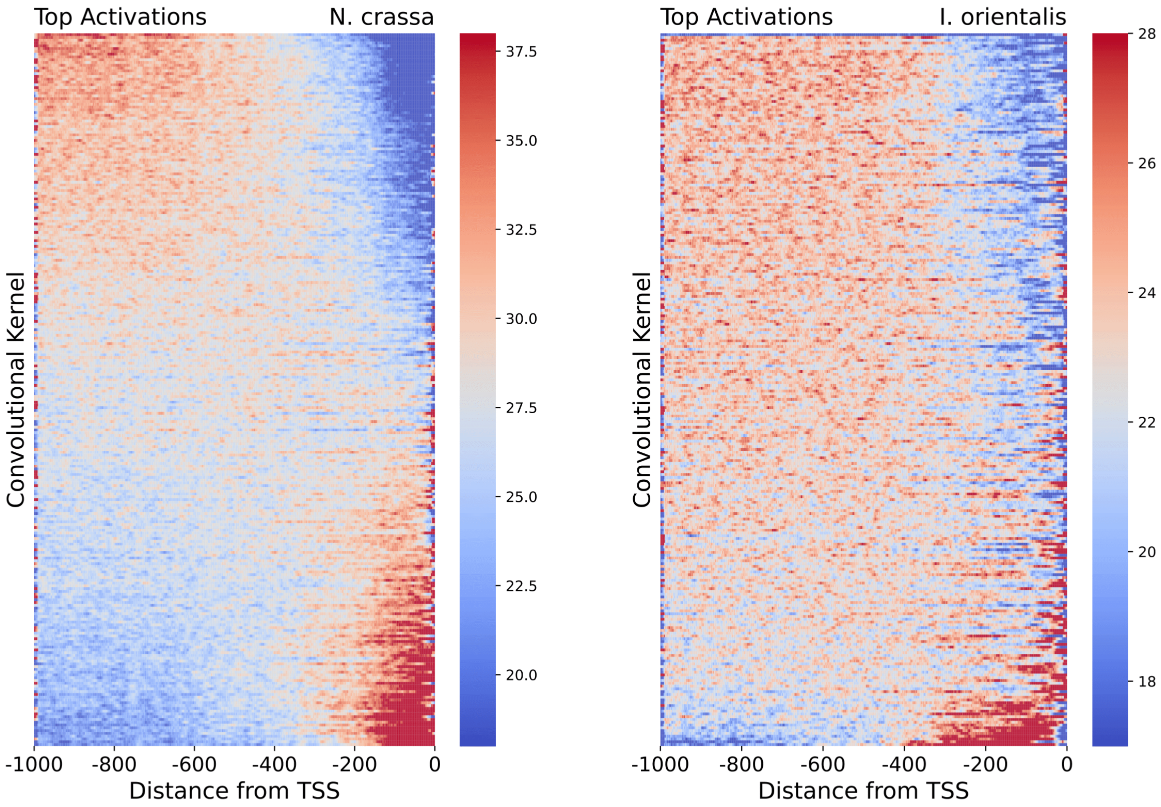


SUPP.FIG. 5. **The heatmaps showing the positional distribution of the top 0.5% of activations for each motif**, i.e. kernel in the first convolutional layer, over all **(left)** *N. crassa* and **(right)** *I. orientalis* promoter sequences. The rows of this heatmap are sorted by the average activation level within 300bp from the transcription start site. Note that most motifs exhibit non-random positional preferences indicative of biological function.

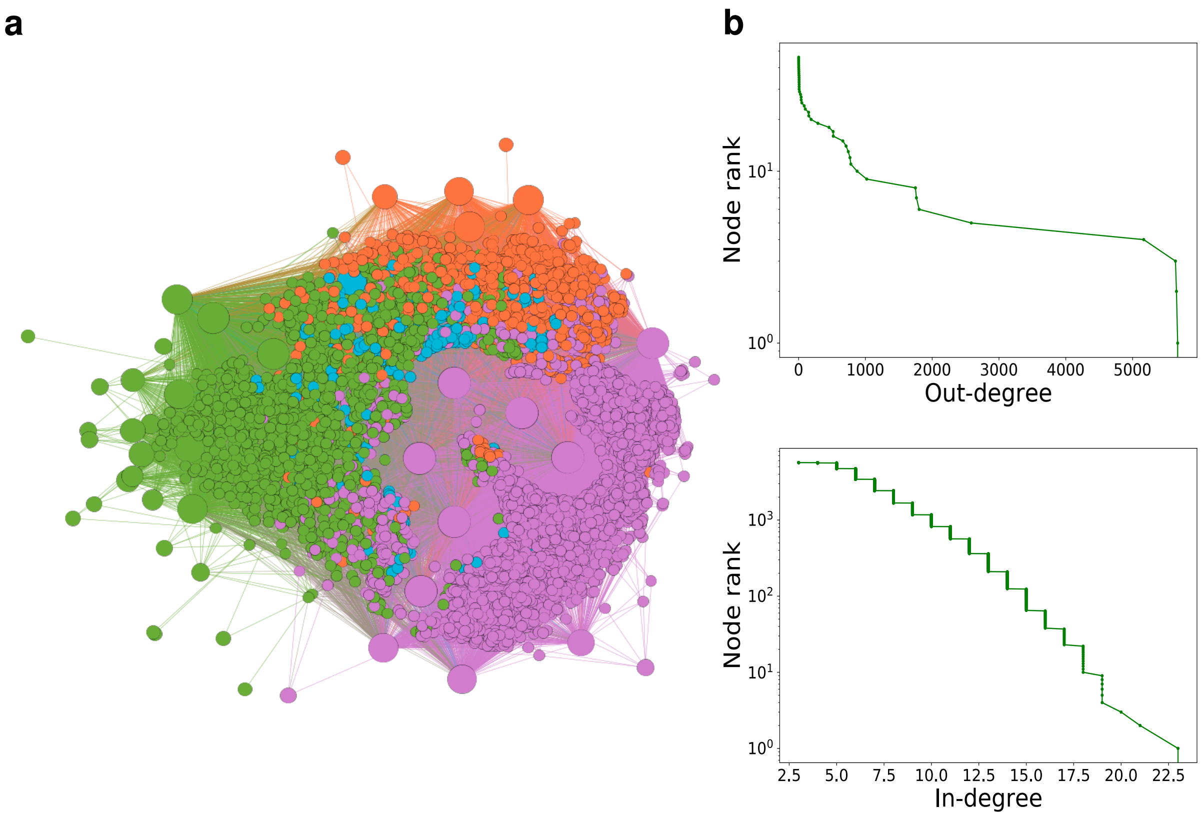


SUPP.FIG. 6. ***N. crassa* Gene Regulatory Network learned by the FUN-PROSE model (a)** The network of TF-target gene interactions obtained by applying a 2.5 standard deviation threshold to the TF-target gene Integrated Gradients scores for the *N. crassa* dataset. Nodes are colored by clusters obtained by modularity optimization. **(b)** Cumulative histogram (number of nodes with degree $>= x$) of out-degrees of TFs and **(c)** in-degrees of target genes.

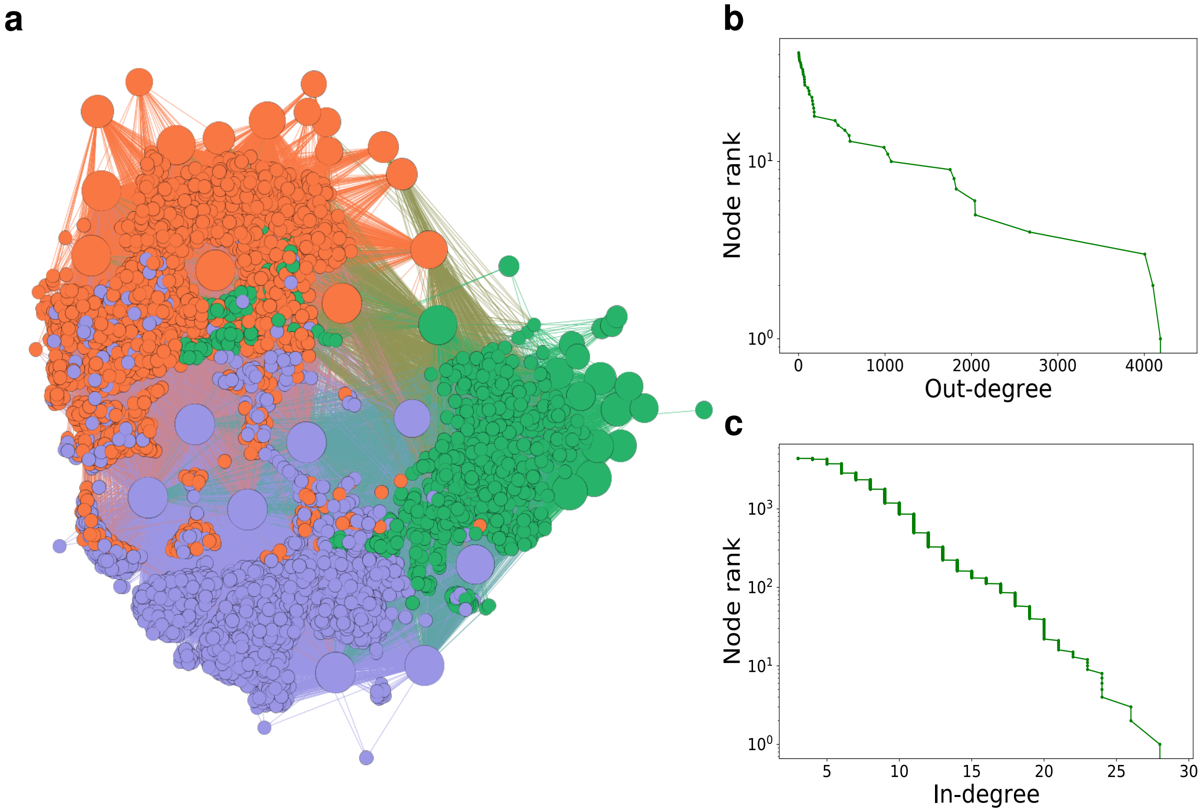


SUPP.FIG. 7. ***I. orientalis*** **Gene Regulatory Network learned by the FUN-PROSE model (a)** The network of TF-target gene interactions obtained by applying a 2 standard deviation threshold to the TF-target gene Integrated Gradients scores for the *I. orientalis* dataset. Red edges mark experimentally confirmed interactions from the YEASTRACT database. **(b)** Cumulative histogram (number of nodes with degree $>= x$) of out-degrees of TFs and **(c)** in-degrees of target genes.
